# Live biotherapeutic throat spray for respiratory virus inhibition and interferon pathway induction

**DOI:** 10.1101/2022.01.25.477549

**Authors:** Irina Spacova, Ilke De Boeck, Eline Cauwenberghs, Lize Delanghe, Peter A. Bron, Tim Henkens, Alix Simons, Imane Gamgami, Leentje Persoons, Ingmar Claes, Marianne F. L. van den Broek, Dominique Schols, Peter Delputte, Samuel Coenen, Veronique Verhoeven, Sarah Lebeer

**Affiliations:** Research Group Environmental Ecology and Applied Microbiology, Department of Bioscience Engineering, University of Antwerp, Belgium; YUN NV, Niel, Belgium; KU Leuven Department of Microbiology, Immunology and Transplantation, Laboratory of Virology and Chemotherapy, Rega Institute, Leuven, Belgium; Laboratory of Microbiology, Parasitology and Hygiene, Department of Biomedical Sciences, University of Antwerp, Belgium; Family Medicine and Population Health (FAMPOP), University of Antwerp, Antwerp, Belgium; Vaccine & Infectious Disease Institute (VAXINFECTIO), University of Antwerp, Antwerp, Belgium

## Abstract

Respiratory viruses such as influenza viruses, respiratory syncytial virus (RSV), and coronaviruses initiate infection at the mucosal surfaces of the upper respiratory tract (URT), where the resident respiratory microbiome has an important gatekeeper function. In contrast to gut-targeting administration of beneficial bacteria against respiratory viral disease, topical URT administration of probiotics is currently underexplored, especially for the prevention and/or treatment of viral infections. Here, we report the selection and formulation of a broad-acting throat spray with live lactobacilli which induce interferon regulatory pathways and are able to inhibit respiratory viruses. Rational selection of *Lactobacillaceae* strains was based on safety, applicability, and potential antiviral and immunostimulatory efficacy in the URT. Three strains, *Lacticaseibacillus casei* AMBR2, *Lacticaseibacillus rhamnosus* GG and *Lactiplantibacillus plantarum* WCFS1 significantly reduced the cytopathogenic effects of RSV, influenza A/H1N1 and B viruses, and HCoV-229E coronavirus in co-culture models with bacteria, virus and host cells. Subsequently, these strains were formulated in a throat spray and human monocytes were employed to confirm the formulation process did not reduce the interferon regulatory pathway-inducing capacity. Administration of the throat spray in healthy volunteers revealed that the lactobacilli were capable of temporary colonization of the throat in a metabolically active form.

## 1. Introduction

Viral respiratory tract infections (RTIs) result in a significant health and economic burden, as recently highlighted by the coronavirus disease 2019 (COVID-19) pandemic [1]. Despite high prevalence of viral RTIs, few prevention or treatment options besides symptom relief are available to primary care patients. Nevertheless, RTIs can have drastic health consequences after the initial infection in the upper respiratory tract (URT), as several viruses such as influenza viruses, respiratory syncytial virus (RSV), human parainfluenza virus (HPIV) and human coronaviruses (*e*.*g*. HCoV-229E and severe acute respiratory syndrome coronavirus 2 (SARS-CoV-2)), can potentially trigger airway tissue disruption, severe inflammation [2,3] and/or subsequent pneumonia [4,5]. Severe disease caused by viruses is often accompanied by reduced type I and III interferon (IFN) production and/or overproduction of pro-inflammatory mediators [3,6,7]. Type I and III IFNs play an important role for innate immunity at mucosal barrier surfaces, such as the respiratory epithelial barrier, where they provide first line antiviral defense mechanisms [8].

The respiratory mucosal surfaces targeted by viruses also harbor a microbiome consisting of resident microorganisms which have important multifactorial gatekeeper functions, including direct inhibition of incoming pathogens, maintaining epithelial barrier function, and immune homeostasis [9]. Importantly, the stability of the core URT microbiome can be compromised during viral RTIs, for instance as observed with influenza [10]. This can facilitate pathobiont overgrowth, as a result of virus-induced damage and immune dysfunction [11,12]. Direct supplementation of beneficial bacteria from the URT microbiome could act on different stages of viral disease, for instance via enhancement of the airway epithelial barrier, direct inhibition of pathogens or stimulation of the immune system, as we have recently reviewed [13]. Stimulation of antiviral immunity is one of the key mechanisms reported in animal models of respiratory viral disease upon topical administration of beneficial bacteria directly to the airways [14,15].

Despite their promise, the number of clinical trials exploring URT microbiome modulation as a strategy against viral RTIs is limited. This is at least in part due to the challenge of selecting appropriate beneficial strains for the URT, and the biotechnical difficulty of formulating live microbes with a long shelf life in a liquid form [13]. Because of the convenient formulation in dry formats, combined with the fact that much more research has been performed on the gut than on the URT microbiome, oral delivery of microbiome therapeutics remains the most common administration route. The oral route focuses on probiotics that can exert systemic antiviral and anti-inflammatory effects via the gut-lung axis [16]. Nevertheless, we have recently isolated probiotic candidates *Lacticaseibacillus casei* AMBR2 [17] and *Streptococcus salivarius* strains [18] capable of respiratory pathogen inhibition from nasopharyngeal samples of healthy humans. Especially *L. casei* AMBR2 has shown potential as a live biotherapeutic product (LBP) due to its capacity to inhibit the growth and inflammatory characteristics of URT pathogens such as *Staphylococcus aureus* and promote epithelial barrier function [19–21]. Furthermore, this strain was able to temporally colonize the URT of all 20 healthy volunteers in a fit-for-purpose nasal spray formulation, without apparent side effects [19]. Also others have shown that the administration of a topical microbiome therapy containing lactic acid bacteria, such as *Lactococcus lactis* W139 [22], a mixture of lactobacilli and *Bifidobacterium* species [23], or commensal *Streptococcus* species [24], in the URT appears safe and could be a better targeted alternative to the indirect gut route. Indeed, lactic acid bacteria have a long history of safe use in other body sites and are therefore also promising for URT applications. Although not as high as reported for other body sites such as the gut and vagina, their prevalence and (relative) abundance in the URT is significant [25]. Moreover, lactic acid bacteria are enriched in healthy individuals compared to individuals suffering from airway diseases such as chronic rhinosinusitis [26] and can have antiviral properties [13]. Implementation of select lactic acid bacteria in high-dose URT-targeting, virus-inhibiting and immune-active formulations is a first prerequisite before clinical studies can be designed.

Based on documented safety, prior human use, annotated genome information and functional properties such as the capacity to inhibit bacterial and fungal pathogens and to modulate immune responses (Table 1), we selected strains of the *Lactobacillaceae* for dedicated immunostimulatory and antiviral screening, resulting in the selection of three strains for formulation in a throat spray. Subsequent application in human volunteers confirmed temporary retention of all three strains in the microbiome.

**Table 1.** Bacterial strains used in this study and their properties.

| Strain | Origin | Properties | References |
| --- | --- | --- | --- |
| ***Lactcaseibacillus casei* AMBR2** | Human URT isolate | URT probiotic candidate with immunomodulatory and anti-pathogenic action against URT pathobionts and the associated inflammation. Has epithelial barrier-promoting properties. Demonstrated safety and colonization upon administration in a nasal spray in healthy volunteers. | [19,21] |
| ***Lactcaseibacillus rhamnosus* GG (ATCC 53103)** | Human gastrointestinal tract isolate | Model oral probiotic strain with anti-pathogenic, barrier-enhancing and immunomodulating properties. Topical URT application in mouse models of influenza led to immune stimulation of antiviral interferon and cytokine pathways. | [14,15,36] |
| ***Lactiplantibacillus plantarum* WCFS1** | Human saliva isolate | Model oral probiotic strain with anti-pathogenic, barrier-enhancing and immunomodulating properties. *In vivo* administration to the URT has not been reported, however NF-κB pathway induction linked to immune tolerance in the duodenum of healthy humans was demonstrated. | [34,47] |
| ***Lactiplantibacillus pentosus* KCA1** | Human vaginal isolate | Well-characterized probiotic strain linked with microbiome modulation upon oral and topical skin application | [48,49] |
| ***Limosilactobacillus reuteri* RC14** | Human vaginal isolate | Vaginal microbiome modulation upon topical application together with *L. rhamnosus* GR-1 | [50] |
| ***Lactcaseibacillus paracasei* Immunitas (DN-114001)** | Origin unclear | Commercial probiotic strain from Actimel Immunitas or Defensis, leads to a decrease in symptoms and duration of respiratory tract infections in clinical trials with children and the elderly | [51,52]<br>Taxonomy cfr. [17] |
| ***Lactcaseibacillus paracasei* Shirota (previously)** | Human gastrointestinal tract isolate | Commercial probiotic strain from the Yakult® product (Yakult Ltd.), ameliorates influenza virus infection in a mouse model | [53]<br>Taxonomy cfr. [17] |
| ***Lactcaseibacillus rhamnosus* B442** | USDA Agriculture Research Service (NRRL) Culture Collection | Proposed probiotic strain for use in ice cream active against enteric pathogens | [54,55] |
| ***Lactcaseibacillus rhamnosus* LC705** | Dairy strain | Commercial probiotic used by companies such as Valio, administered as part of probiotic mixtures in yogurt | [56] |
| ***Lactiplantibacillus plantarum* ATCC8014** | Deposited to ATCC by E. McCoy as “17-5 (*Lactobacillus arabinosus*)” | Model strain of *L. plantarum* with anti-pathogenic properties, | [57] |
| ***Lactobacillus acidophilus* LMG 8151** | Isolate from commercial acidophilus milk | Commercial probiotic strain, deposited by the Culture Collection University of Göteborg (CCUG), Department of Clinical Bacteriology, Institute of Clinical Bacteriology, Immunology, and Virology under number CCUG 12853 | [58] |
| ***Latilactobacillus sakei* AMBR8** | Healthy human URT isolate | Strain isolated from and potentially adapted to the human URT (e.g., adherence to airway epithelial cells) with immunomodulatory and anti-pathogenic action against URT pathobionts | [19] |
| ***Lactiplantibacillus plantarum* AMBR9** | Healthy human URT isolate | Strain isolated from and potentially adapted to the human URT with anti-pathogenic action against URT pathobionts | [19] |
| ***Escherichia coli* DH5α** | Laboratory strain | Laboratory strain of *E. coli* carrying mutations that facilitate efficient transformation | [59] |

## 2. Results

### 2.1. Screening for interferon regulatory pathway- and NF-κB-inducing *Lactobacillaceae* strains

A selection of strains described in Table 1 that have previously been effectively used as probiotics in clinical trials (*e*.*g*., *Lacticaseibacillus rhamnosus* GG, *Lactiplantibacillus plantarum* WCFS1, *Limosilactobacillus reuteri* RC-14) and/or isolated from the URT and previously demonstrated to interact with URT cells (*e*.*g. L. casei* AMBR2, *Lactiplantibacillus plantarum* AMBR9) were used in this first broad screening. The strains were evaluated for their capacity to activate the interferon regulatory factor (IRF) and nuclear factor kappa B (NF-κB) immune signaling pathways in human THP-1 Dual monocytes involved in antiviral responses. Of all tested strains, *L. plantarum* WCFS1 most strongly activated the IRF pathway (Figure 1A), as well as NF-κB (Figure 1B). IRF induction was shown to be strain-specific with for example *L. rhamnosus* GG stimulating a stronger IRF and NF-κB activation compared to the other *L. rhamnosus* strains tested.

**Fig. 1.**
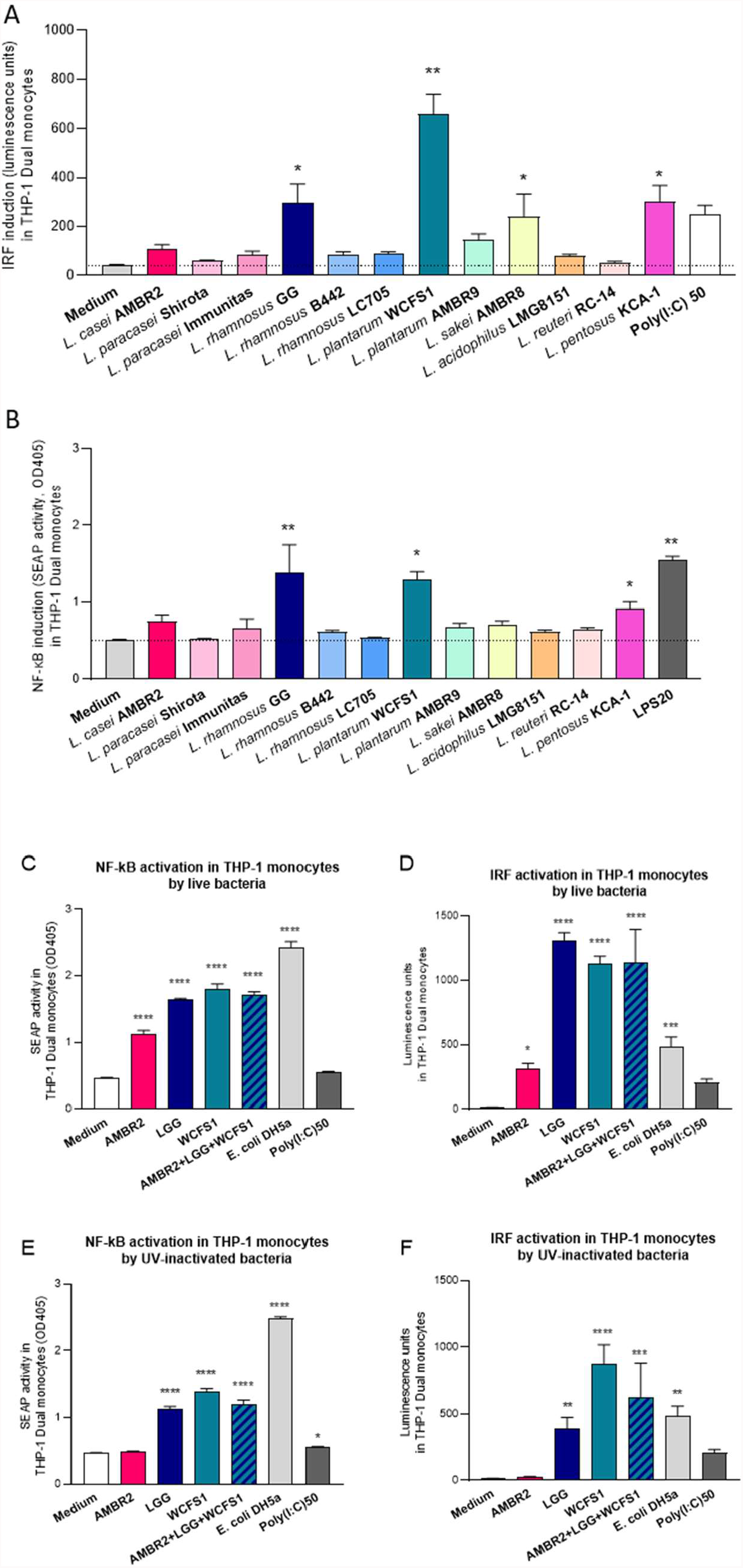
(A) Stimulation of interferon regulatory factors (IRFs) and (B) nuclear factor (NF)-κB by selected *Lactobacillaceae* strains in human monocytes, and (C-G) *L. casei* AMBR2, *L. rhamnosus* GG, *L. plantarum* WCFS1 and their combination AMBR2/WCFS1/LGG activate immune pathways involved in antiviral responses. Live *L. casei* AMBR2, *L. rhamnosus* GG and *L. plantarum* WCFS1 and their combination induce (C) nuclear factor (NF)-κB and (D) interferon regulatory factors (IRFs) in human THP-1 Dual monocytes upon co-incubation. UV-inactivated *L. rhamnosus* GG and *L. plantarum* WCFS1 and their combination with *L. casei* AMBR2 also induce (E) NF-κB and (F) IRFs in human THP-1 Dual monocytes. The medium condition represents the cells as such and serves as a baseline, while Poly(I:C) at 50 μg/ml with Lipofectamine (Poly(I:C) 50) serves as control IRF inducer and LPS at 20 ng/ml (LPS20) and LPS-producing *E. coli* DH5 serve as control NF-κB inducer. Data is depicted as mean±SD per condition. *p<0.05, **p<0.01, ***p<0.001 and ****p<0.0001 as determined by a One-way ANOVA test followed by Dunnett’s multiple comparisons test compared to the medium condition.

Based on these data and previous knowledge on their activity, *L. casei* AMBR2, *L. plantarum* WCFS1 and *L. rhamnosus* GG were selected for further analyses, as these strains have additional strain-specific properties beneficial in the URT (Table 1). Hence, a product containing a combination of strains could be more efficient due to multifactorial action, providing that the strains do not inhibit each other’s activity. Indeed, combining the three strains in equal ratios also resulted in significant induction of IRF and NF-κB pathways in human monocytes, comparable to the levels observed when *L. rhamnosus* GG and *L. plantarum* WCFS1 were used separately (Figure 2C-F). Of note, the NF-κB stimulation by lactobacilli was significantly lower compared to the non-probiotic LPS-containing Gram-negative control *Escherichia coli* DH5α (Figure 1C, E). To assess whether the bacteria had to be metabolically active to induce immunostimulatory effects, ultraviolet (UV)-inactivated strains were used (Figure 1E-F). Both UV-inactivated *L. plantarum* WCFS1 and *L. rhamnosus* GG, as well as the mix of the three strains, still significantly induced the IRF and NF-κB pathways, although to a lesser extent than their viable counterparts (Figure 1E-F).

**Fig. 2.**
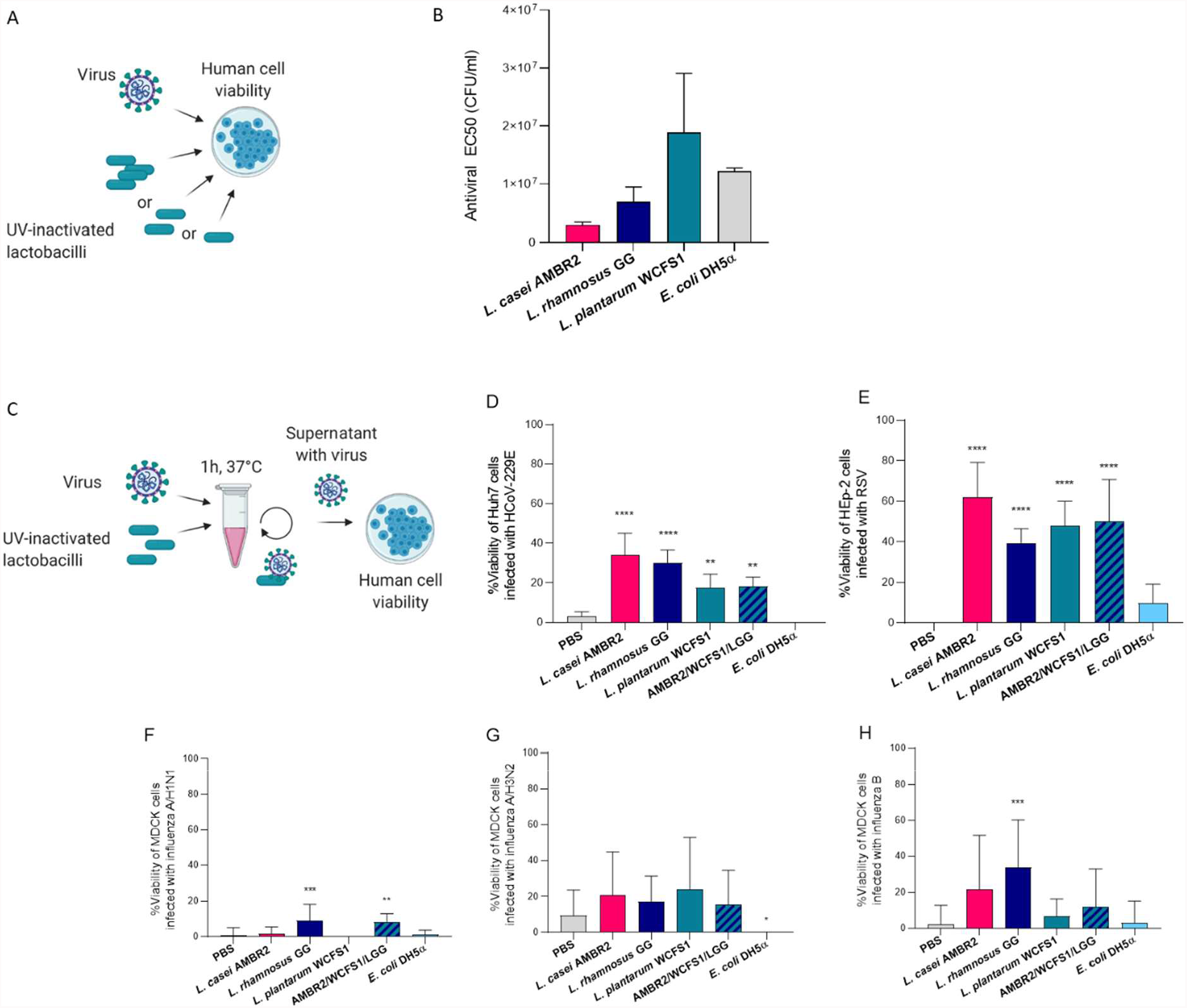
*L. casei* AMBR2, *L. rhamnosus* GG, *L. plantarum* WCFS1 and their combination AMBR2/WCFS1/LGG decrease cytopathogenic effects of HCoV-229E, RSV and influenza A/H1N1, A/H3N2 and B viruses *in vitro*. *L. casei* AMBR2, *L. rhamnosus* GG and *L. plantarum* WCFS1 that were added together with the HCoV-229E virus (A, B) or pre-incubated (C) with HCoV-229E (D), RSV (E), influenza A/H1N1 (F), A/H3N2 (G) or influenza B (FH could inhibit the cytopathic effects induced in human cells. EC50: 50% Effective concentration producing 50% inhibition of virus-induced cytopathic effects, as determined by measuring the cell viability with the colorimetric MTS cell viability assay. CFU: colony-forming units. PBS serves as control that does not affect viral infection, *E. coli* DH5α represents a non-probiotic laboratory strain. Data depicted as mean±SD per condition. *p<0.05, **p<0.01, ***p<0.001 and ****p<0.0001 as determined by a One-way ANOVA test followed by Dunnett’s multiple comparisons test compared to the medium or PBS conditions.

### 2.2. Direct antiviral activity of selected *Lactobacillaceae* strains against respiratory viruses

Probiotic screening for direct antiviral activity is not routinely implemented because of the complexity of working with tripartite systems consisting of bacteria, viruses and host cells. Here, we implemented such innovative assays for common respiratory viruses. The inhibitory effects of the three selected *Lactobacillaceae* strains were evaluated against RSV, influenza A/H3N2, A/H1N1 and B viruses, and human coronavirus strain 229 (HCoV-229E), all representing viruses that are a common cause of RTIs [27]. HCoV-229E generally causes mild symptoms, but it has several features in common with SARS-CoV-2, such as homologous epitopes of the spike protein, and is suitable for high throughput screening due to biosafety level 2 [28]. To avoid bacterial metabolites interfering with the assay read-outs, the lactobacilli were UV-inactivated for the assays, retaining the capacity of bacterial cell surface molecules to sequester or block viral particles [29]. First, we established the concentration of bacteria that had to be simultaneously added with the HCoV-229E virus to inhibit the virus-induced cytopathic effect in susceptible human Huh7 cells by 50% (antiviral EC50) (Figure 2A,B). *L. casei* AMBR2 and *L. rhamnosus* GG showed the strongest effect, with respective concentrations of 2.57×10^6^ colony-forming units (CFU)/ml and 5.06×10^6^ CFU/ml required for 50% viral inhibition, which was lower than the required concentration of 1.26×10^7^ CFU/mL for the non-probiotic *E. coli* DH5α strain. For *L. plantarum* WCFS1, the required concentration for 50% viral inhibition was 1.17×10^7^ CFU/ml.

Subsequently, we also evaluated a set-up where the three bacterial strains or their combination in equal ratios were pre-incubated with HCoV-229E, RSV or influenza viruses, allowing trapping and/or inactivation of virus particles, and afterwards unbound virus particles were added to human cells (Figure 2C). HCoV-229E (Figure 2D) and RSV (Figure 2E) pre-incubated with *L. casei* AMBR2, *L. rhamnosus* GG, *L. plantarum* WCFS1 or their combination significantly lost the capacity to reduce human cell viability compared to virus pre-incubation with phosphate-buffered saline (PBS) which served as an inactive control. For example, the percentage of viable Huh7 cells infected with HCoV-229E was on average 2.9% for the PBS condition, and 34%, 30%, and 18% for *L. casei* AMBR2, *L. rhamnosus* GG, and *L. plantarum* WCFS1 conditions, respectively. Using the combination of the three strains also resulted in 18% of viable Huh7 cells, similar to *L. plantarum* WCFS1 only. Pre-incubation with *L. rhamnosus* GG or a combination of the three lactobacilli also significantly reduced the cytopathic effects induced by the common influenza A/H1N1 virus compared to PBS (Figure 2F). A reduction of cytopathic effects of influenza B was also observed after pre-incubation of *L. rhamnosus* GG (Figure 2H), while the effects of the tested lactobacilli on the cytopathic effects of influenza A/H3N2 virus were less pronounced (Figure 2G). The pre-incubation of the non-probiotic control *E. coli* DH5α with the tested viruses did not improve viability of human cells, while the cell viability in all lactobacilli conditions was significantly higher compared to *E. coli* DH5α. The potential antiviral activity of *L. casei* AMBR2, *L. rhamnosus* GG, and *L. plantarum* WCFS1, in addition to their other beneficial properties (Table 1) and interferon pathway induction capacity (Figure 1), led us to evaluate their combination in a throat spray formulation.

### 2.3. Formulation of viable *Lactobacillaceae* strains in a throat spray

*L. casei* AMBR2, *L. rhamnosus* GG, and *L. plantarum* WCFS1 were formulated into an oral/throat-targeting spray, based on a combination of bacterial powders in an oil suspension. Although most probiotic sprays for the URT currently available consist of a bacterial suspension in saline or PBS [30], oil was chosen to increase bacterial retention in the throat and to ensure a sufficient dosage of viable probiotic CFU counts. First, the viability of each bacterial strain in freeze-dried powder form was evaluated at 4°C and 25°C (Figure 3A). For all three strains, the viability at 4°C remained stable over time. For *L. casei* AMBR2 and *L. plantarum* WCFS1, no log reductions were observed, with 2.79 × 10^10^ CFU/g powder and 3.56 × 10^11^ at start for AMBR2 and WCFS1, respectively, to 2.21 × 10^10^ CFU/g powder and 3.15 × 10^11^ CFU/g powder after 26 weeks. For *L. rhamnosus* GG, the viability decreased from 1.18 × 10^11^ CFU/g powder to 7.15 × 10^10^ CFU/g. At room temperature (25°C), *L. plantarum* WCFS1 seemed most stable, with 1 log reduction at 26 week (5.84 × 10^10^), while this was 8.57 × 10^9^ for *L. rhamnosus* GG and 2.02 × 10^7^ for *L. casei* AMBR2.

**Fig. 3.**
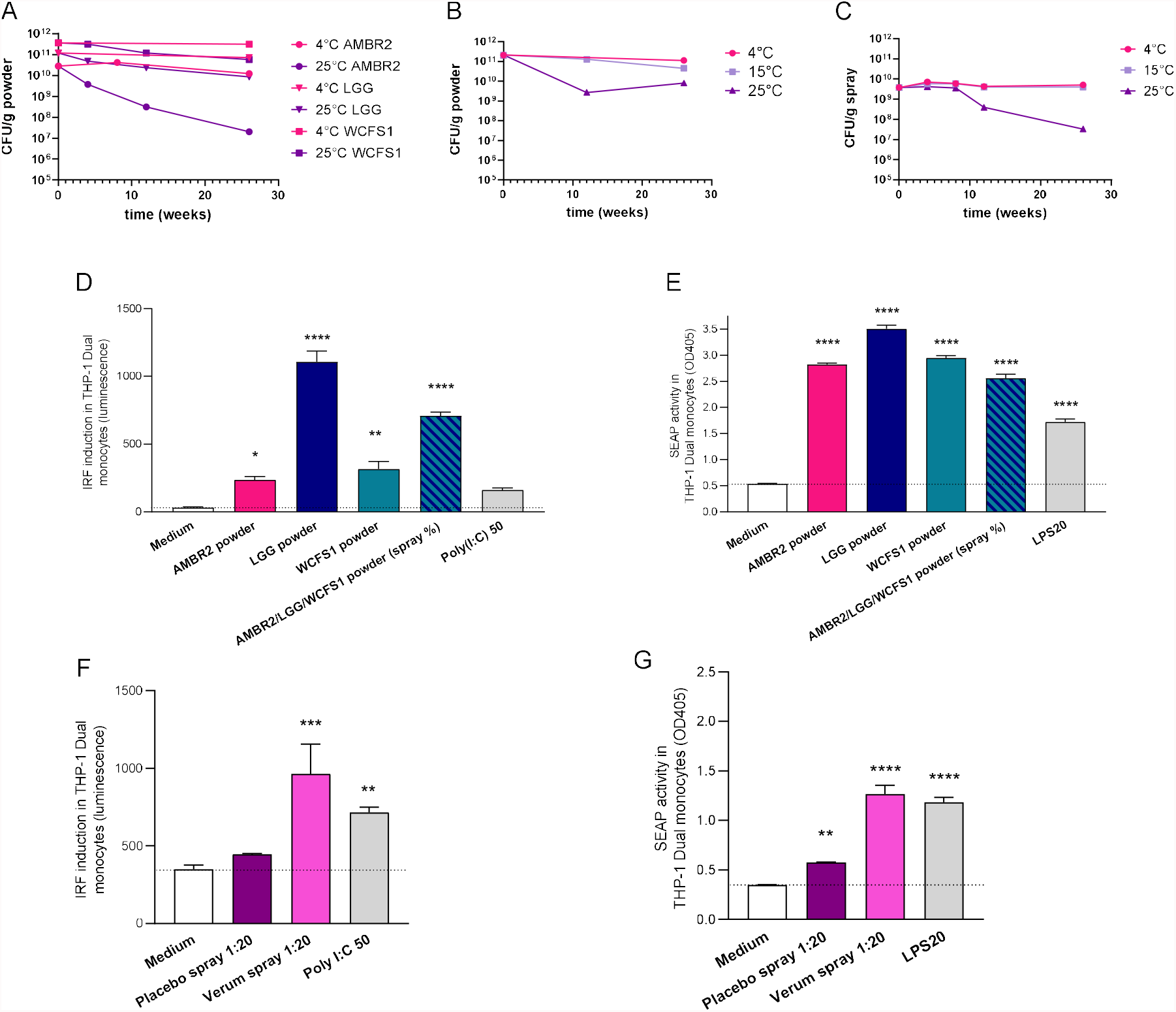
Viability of individual and combined *L. casei* AMBR2, *L. plantarum* WCFS1 and *L. rhamnosus* GG in (A-B) powder or (C) the throat spray formulation; Immunostimulatory activity of the powders (D-E) and the placebo and verum spray formulation without or with lactobacilli, respectively (F-G). The viability of the strains combination was evaluated at different storage temperatures over time; 4°C, 15°C and 25°C. CFU: colony-forming units. The medium condition represents the cells as such and serves as a baseline, Poly(I:C) at 50 μg/ml with Lipofectamine (Poly(I:C)50/Lipo) serves as control IRF inducer and LPS at 20 ng/ml (LPS20) serves as control NF-κB inducer. Data is depicted as mean ± SD per condition. *p<0.05, **p<0.01, ***p<0.001 and ****p<0.0001 as determined by a One-way ANOVA test followed by Dunnett’s multiple comparisons test compared to the medium condition (dotted line).

We next evaluated different concentrations of the strains in the mixture. *L. casei* AMBR2 at 50%, *L. plantarum* WCFS1 at 33.3%, and *L. rhamnosus* GG at 16.7% were found the optimal ratio upon long-term storage at room temperature, which reflected the intended storage conditions. Next, the viability of the combined bacterial strains in powder form (Figure 3B) and in the throat spray formulation (Figure 3C) was evaluated at 4°C, 15°C and 25°C. For the mixed powders (Figure 4B), viability decreased slightly from 2.09 × 10^11^ CFU/g at the start, to 1.11 × 10^11^, 4.51 × 10^10^ and 8.01 × 10^9^ CFU/g at 26 weeks of storage at 4°C, 15°C and 25°C, respectively. For the spray formulation (Figure 3C), viability starting with 3.78 × 10^9^ CFU/g spray remained stable at 4°C and 15°C at 26 weeks. At 25°C, a 2 log reduction was observed (3.3 × 10^7^ CFU/g) at 26 weeks.

**Fig. 4.**
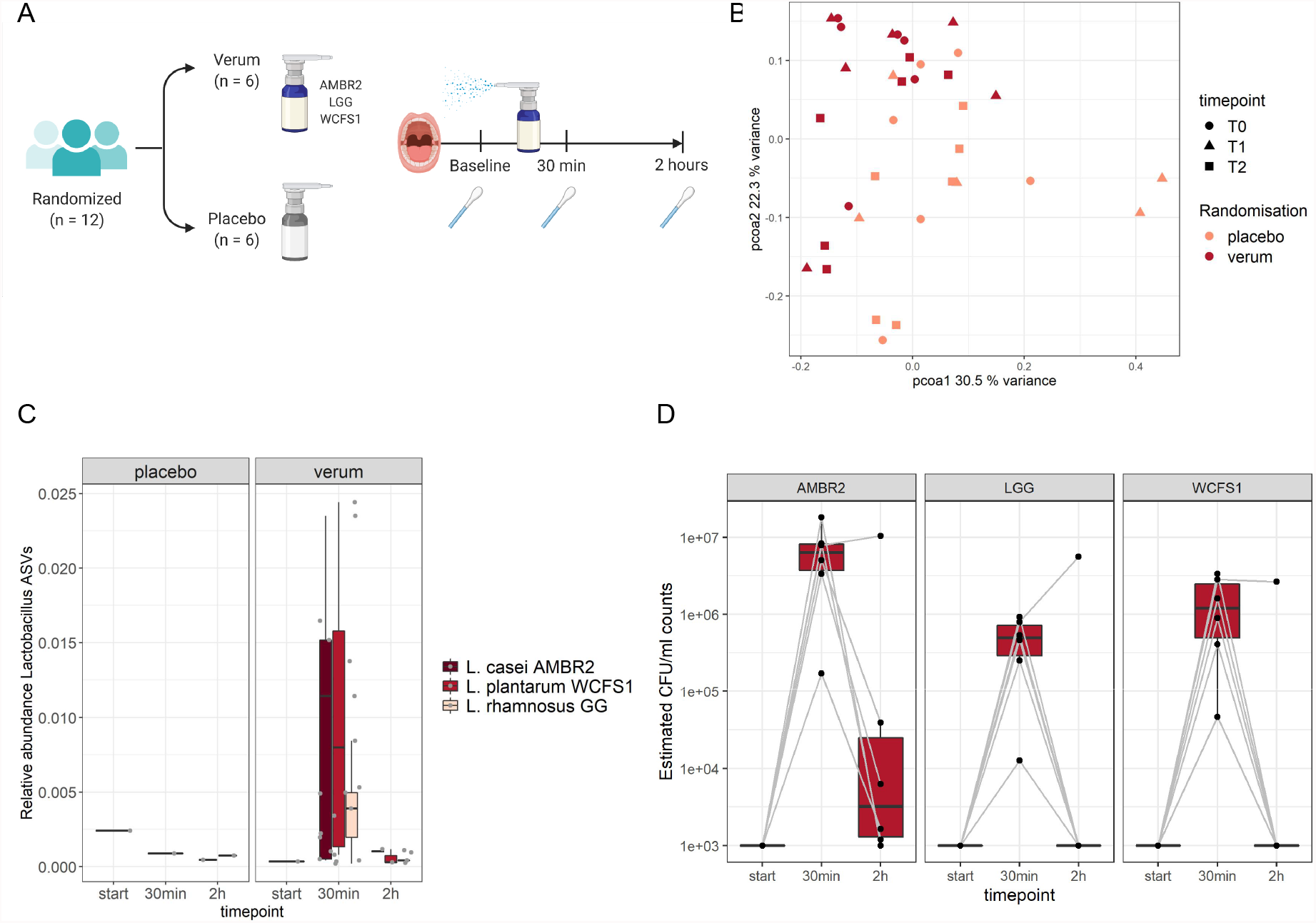
Evaluation of lactobacilli retention within the throat microbiome of healthy volunteers after spray application. (A) Study set-up; (B) PCoA plot of throat microbiome data based on microbiome analysis via 16S rRNA amplicon sequencing; (C) Relative abundance of the administered lactobacilli in throat swabs based on microbiome analysis via 16S rRNA amplicon sequencing; (D) Relative abundance of the administered lactobacilli in throat swabs based on qPCR analysis. Throat swabs were collected at baseline (T0), and 30 minutes (T1) and 2 hours (T2) after the throat spray was used. The presence of *L. casei* AMBR2, *L. rhamnosus* GG, and *L. plantarum* WCFS1 was evaluated via 16S rRNA amplicon sequencing (relative abundances) in panel C. At 30 minutes and 2 hours, qPCR with species-specific primers was used to estimate the CFU/ml counts in the verum group in panel D. Based on the standard curve, the detection limit was estimated to be at 10^3^ CFU/ml.

We subsequently confirmed the retention of immunostimulatory activity in human monocytes of the strains and their combination in powder form (Figure 3D-E), and in the spray formulation in oil (Figure 3F-G). All single strains in powder form and their combination were still capable of significant IRF and NF-κB induction (Figure 4D-E) at a dose of 10^8^ CFU/ml, which corresponds to the *L. casei* AMBR2 concentration per puff previously tested in healthy volunteers [19]. The immunostimulatory action of the throat spray formulation with the three strains in an oil suspension was also compared to a placebo oil formulation without lactobacilli. The throat spray formulation with *L. casei* AMBR2 at 50%, *L. plantarum* WCFS1 at 33.3%, and *L. rhamnosus* GG at 16.7% in oil significantly induced IRF and NF-κB in human monocytes (Figure 4F-G). While the placebo formulation also induced NF-κB, albeit to a lower degree compared to the spray formulation with lactobacilli, it did not significantly affect IRF.

### 2.4. Evaluation of throat colonization by lactobacilli in healthy volunteers

Finally, we evaluated the retention of the lactobacilli in the formulated spray in the throat of 12 healthy volunteers via cultivation, quantitative polymerase chain reaction (qPCR) and 16S rRNA amplicon sequencing using a longitudinal placebo-controlled sampling set-up (Figure 4A). The volunteers used the verum spray at the start of the study by spraying two puffs containing approximately 9.5×10^8^ CFU of lactobacilli, or the placebo spray not containing lactobacilli, and throat swabs were collected at baseline, after 30 min, and after 2 hours (Figure 4A).

Microbiome analysis of the throat swabs showed that the dominant bacterial genera in the throat belong to canonical throat commensals, such as *Prevotella, Veillonella*, and *Streptococcus* species. A principal coordinate analysis (PCoA) plot based on the microbiome data across all time points did not reveal a clear clustering per treatment group at any of the tested time points (Figure 4B). A clear difference in relative abundances of *Lactobacillaceae* amplicon sequence variants (ASVs) was observed between the placebo and verum spray groups, especially 30 minutes after the spray was used (Figure 4C). Three *Lactobacillus* ASVs, corresponding to the administered strains, were detected in the verum group 30 minutes after spray administration. *Lactobacillus* ASV 1 (*L. rhamnosus*) was detected in all 6 participants in the verum group, while *Lactobacillus* ASV 3 (*L. casei*) and *Lactobacillus* ASV 7 (*L. plantarum*) were detected in 5 out of 6 participants. In the placebo group, these *Lactobacillus* ASV were not detected, except for one participant that had low endogenous relative abundances of the *Lactobacillus casei* ASV after 30 minutes. After 2 hours, 5 out of 6 participants in the verum group still had detectable *Lactobacillus* ASVs.

To confirm and quantify the high abundances of the administered strains observed by sequencing DNA derived from samples in the verum group after bacterial administration, we aimed to estimate the CFU/ml counts based on targeted qPCR (Figure 4D). In line with the sequencing data, after 30 minutes, the estimated CFU counts for *L. rhamnosus* GG in the verum group were between 1.26 × 10^4^-9.24 × 10^5^ CFU/ml. For *L. casei* AMBR2, estimated CFU/ml counts ranged from 1.72 × 10^5^ to 1.8 × 10^7^ CFU/ml and for *L. plantarum* WCFS1 from 4.63 × 10^4^ CFU – 3.36 × 10^6^ CFU. After 2 hours, the amount of detected lactobacilli decreased. *L. rhamnosus* GG and *L. plantarum* WCFS1 were not detected anymore except in one participant. *L. casei* AMBR2 on the other hand, which was administered in the highest ratio of 50% in the spray, was still detected in 5 of the 6 participants, with a median CFU/ml count of 4×10^3^ CFU/ml.

In addition to analyzing the DNA of the bacteria, we also cultivated throat swabs to evaluate whether the administered lactobacilli were still viable. Cultured throat swabs from the verum group demonstrated colony morphologies typical for the three administered *Lactobacilaceae* strains, and the identity was confirmed via colony PCR and sequencing of the *16S rRNA* gene, confirming that the administered strains are not only detected in the throat via their DNA, but also remain viable.

## 3. Discussion

Topical microbiome therapeutics can act on different stages of viral infection by multiple modes of action, which can potentially prevent a broad range of viral diseases [13]. Here, we designed and implemented a dedicated pipeline to select lactobacilli for a topical throat spray against viral RTIs (Figure 5A). We demonstrated the immunostimulatory and antiviral effects of selected lactobacilli, and also considered their safety, applicability and previously reported beneficial properties such as barrier enhancement and antipathogenic effects, important during viral RTIs. Starting from this screening, we formulated *L. casei* AMBR2, *L. rhamnosus* GG, and *L. plantarum* WCFS1 in a microbiome throat spray, confirmed their immunostimulatory activity in the spray formulation and the temporary colonization of the administered strains in throats of healthy volunteers.

**Fig. 5.**
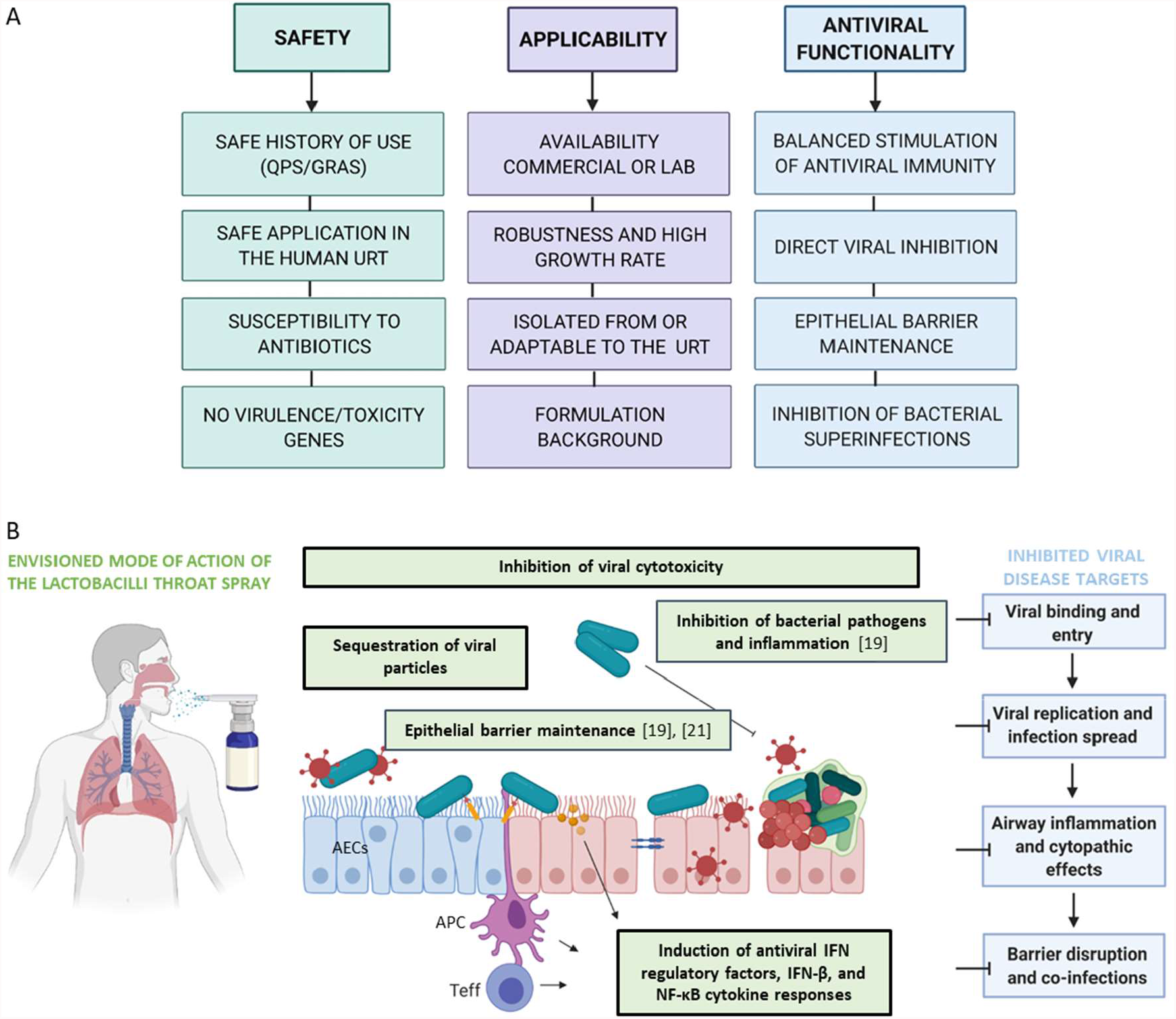
(A) Rationale for selection of *Lactobacillaceae* strains for topical application against respiratory viral disease. (B) Documented mechanisms from this and previous studies through which select beneficial lactobacilli formulated as an URT spray can promote the antiviral activity of the URT microbiome. The modes of action explored in this study are shown in bold frames, while available data from previous studies on *L. casei* AMBR2, *L. rhamnosus* GG and *L. plantarum* WCFS1 are indicated in non-bold green boxes [19,21]. IFN: interferon; AECs: airway epithelial cells; APC: antigen-presenting cell; Teff: T effector cell (based on [13]).

An important property of the selected *Lactobacilaceae* strains was the activation of IFN regulatory pathways and NF-κB that could promote antiviral immune defenses upon topical application of the selected lactobacilli in the URT, based on the documented key role of these pathways [31]. Our results highlight that different lactobacilli isolates have strikingly variable capacities for NF-κB and IFN regulatory pathway stimulation, both at the species and strain level, highlighting that thorough screening is required to select the optimal strain for each application. Specifically, we have identified *L. plantarum* WCFS1 and *L. rhamnosus* GG as two promising strains leading to significant IFN regulatory pathway induction both in viable and, to a lesser degree, UV-inactivated form. This indicates their highest potential as live probiotics, but also as postbiotics because of retained activity in inanimate forms [32]. The strong induction of IFN pathway by *L. plantarum* WCFS1 is in line with earlier observations demonstrating especially strong activation by this strain of the antiviral IFIT1 (Interferon Induced Protein With Tetratricopeptide Repeats 1) gene in differentiated human monocytes [33]. Similarly, intranasal supplementation with *L. rhamnosus* GG was previously shown to increase survival in mouse models of influenza infection by stimulating transcription of protective type I IFN genes in the neonatal airways [15]. These immunostimulatory effects could be extrapolated to humans, as in the duodenum of healthy participants *L. plantarum* WCFS1 induced the NF-κB pathway linked to immune tolerance [34], while consumption of *L. rhamnosus* GG up-regulated IFN-induced genes and cytokine-encoding genes [35].

Importantly, we demonstrated that the combination of the selected lactobacilli retained their immunostimulatory capacity not only when cultured in laboratory conditions, but also as part of the final throat spray formulation. These results are in line with our observation that the viability of the *L. casei* AMBR2, *L. rhamnosus* GG, and *L. plantarum* WCFS1 is maintained after freeze-drying of the bacteria and resuspension in oil. The use of oil in the final spray formulation, in combination with the established adherence capacity to the respiratory epithelium of the bacterial strains themselves [19,36,37] ensures their temporary retention in the throat. To the best of our knowledge, other oral sprays with probiotics have not yet used oil, but instead suspended the bacteria in saline [24]. In this study, retention of the administered lactobacilli was confirmed *in vivo* in healthy volunteers. We found that 30 minutes after spray administration, live *L. casei* AMBR2, *L. rhamnosus* GG, and *L. plantarum* WCFS1 and their DNA were detected in the throat. A contact time of 30 minutes has previously been shown to be sufficient for immune response initiation *in vitro*, including NF-κB induction by bacterial lipopolysaccharides in RAW 264.7 macrophages [38] and in human cell lines by interleukin 1β and tumor necrosis factor alpha [39]. The spray was successfully tested in healthy volunteers, which opens possibilities for its preventive use. However, the dynamics of bacterial colonization in a disturbed throat environment during an active viral RTI could be different, therefore the spray should also be tested in volunteers with viral RTIs to better assess its therapeutic potential. Promisingly, our experiments here already established the delivery of a dose of live cells at their target site in healthy volunteers comparable to the amount of cells that was capable to counteract viral infection *in vitro*.

Our formulation ensures that the throat is the main target for bacterial interaction, in contrast to other studies on oral bacteriotherapy where the formulation is not tailored to URT applications, with most of the bacteria ending up in the gut [40]. This is important, because studies in animal models show that applying immunomodulatory lactobacilli at the site of respiratory infection or inflammation rather than in the gastrointestinal tract, promotes a more direct and efficient contact with specific cells of the URT immune system [41–44].

In addition to the important immunostimulatory effects on host cells, our *in vitro* results also demonstrate direct inhibitory activity of *L. casei* AMBR2, *L. rhamnosus* GG and *L. plantarum* WCFS1 on HCoV-229E, RSV and influenza. While production of L-and D-lactic acid has previously been shown to reduce influenza A virus A/WSN/33 (H1N1) titers *in vitro* [45], reduction of viral cytopathogenic effects upon direct contact with metabolically inactive lactobacilli has not yet been explored in detail, especially not for promising URT isolates such as *L. casei* AMBR2. Since the lactic acid bacterium *Enterococcus faecium* NCIMB 10415 is capable of direct adsorptive trapping of swine influenza H1N1 and H3N2 viral particles [46], we hypothesize that *L. casei* AMBR2, *L. rhamnosus* GG and *L. plantarum* WCFS1 are also capable of sequestering HCoV-229E, RSV and influenza viral particles and directly reducing their infectivity and transmission.

In combination with previous results, our comprehensive *in vitro* testing substantiated the selection of *L. casei* AMBR2, *L. rhamnosus* GG and *L. plantarum* WCFS1 as three promising strains capable of broad activity against different key aspects of viral respiratory diseases when combined in a URT spray (Figure 5B). In our previous research, the potential of *L. casei* AMBR2 as topical probiotic for the respiratory tract is already well-established [20,21,25]. An important feature of this strain in the context of viral disease is its strong ability to enhance the airway epithelial barrier, as well as its adaptation to the unique URT environment, including catalase genes and adherence to the airway epithelium [17,20,21,25]. Probiotics such as *L. casei* AMBR2 and *L. rhamnosus* GG also inhibit URT pathobionts that can increase the morbidity and mortality of viral respiratory diseases, including *Haemophilus influenzae, S. aureus* and *Moraxella catarrhalis* [12,19,36]. These features, in combination with the newly reported antiviral activity in this study pave the way for a more directed topical use of the developed *L. casei* AMBR2, *L. rhamnosus* GG and *L. plantarum* WCFS1 throat spray against respiratory viral disease.

## 4. Conclusions

The development of rationally-selected, evidence-based sprays with probiotic lactobacilli and their employment in clinical trials is long overdue, especially in the context of viral RTI. Based on their safety, applicability in the airways, antiviral action and stimulation of antiviral immunity, *L. casei* AMBR2, *L. rhamnosus* GG and *L. plantarum* WCFS1 were selected as promising strains, followed by optimal formulation for topical application in the URT in a throat spray. Temporary throat colonization by the administered lactobacilli was demonstrated. The spray is a stepping stone for prevention and treatment of viral respiratory tract infections with locally applied beneficial bacteria, and will be further evaluated in primary care patients with viral respiratory tract infections.

## 5. Material and Methods

### 5.1. Bacterial strains and culture

Bacterial strains used in this study and their properties are listed in Table 1. *Lactobacillaceae* strains were grown statically at 37°C in de Man, Rogosa and Sharpe (MRS) broth (Difco, Erembodegem, Belgium). *Escherichia coli* DH5α was cultured aerobically at 37°C in Luria-Bertani (LB) broth. For co-incubation with viruses or human cells, bacterial pellets were obtained by centrifugation at 2000 g for 10 min and washed in sterile phosphate-buffered saline (PBS). After resuspension in corresponding media (for experiments with human cells) or PBS (for antiviral activity experiments), UV inactivation of bacteria was achieved in a biosafety level 2 cabinet using four 15 min rounds of UV irradiation followed by vortexing. Bacteria were plated out to confirm inactivation.

### 5.2. NF-κB and IRF induction in THP1-Dual monocytes

THP1-Dual monocytes (Invivogen) were maintained in RPMI 1640 (ThermoFisher Scientific) medium with 10% Fetal Calf Serum (FCS), 25 mM HEPES and 2 mM L-glutamine at 37°C, 5% CO_2_. For experiments with bacteria, THP1-Dual cells were seeded in a 96-well plate at a concentration of 10^5^ cells/well. Bacteria were added to the cells at 10^6^ CFU/well for live bacteria from cultures, 10^7^ CFU/well for UV-inactivated bacteria from cultures, and 10^8^ CFU/well for powdered bacteria. Spray was added at a 1:20 dilution. The plate was incubated for 24 hours at 37°C and 5% CO_2_. Induction of NF-κB was assessed based on SEAP reporter activity at 405 nm with the Synergy HTX Plate Reader (BioTek) after the addition of a para-Nitrophenylphosphate (pNPP) buffer. Induction of IRF was assessed based on luciferase reporter luminescence activity with the Synergy HTX Plate Reader (BioTek) after the addition of the QUANTI-Luc™ (InvivoGen) buffer. Poly (I:C) with Lipofectamine 2000 (Invitrogen) at 50 μg/ml for IRF induction or lipopolysaccharides (LPS) from *E. coli* (Sigma) at 20 ng/mL for NF-κB induction were used as positive controls.

### 5.3. Cell lines and viruses for antiviral assays

Human liver carcinoma cell line Huh7 (CLS Cell Lines Service, Germany), human epidermoid carcinoma HEp-2 (ATCC CCL-23) and Madin-Darby canine kidney (MDCK) cells, a gift from Dr. M. Matrosovich (Marburg, Germany), were maintained in Dulbecco’s Modified Eagle Medium (DMEM, Gibco Life Technologies) supplemented with 8% heat-inactivated fetal bovine serum (HyClone, GE Healthcare Life Sciences), and maintained at 37°C under 5% CO_2_.

Human coronavirus (HCoV-229E) and respiratory syncytial virus (RSV Long) were purchased from ATCC (VR-740 and VR-26, respectively). The human influenza virus strains used are as follows: A/Ned/378/05 (A/H1N1 subtype) and B/Ned/537/05 (B/Yamagata lineage), clinical isolates generously donated by Prof. R. Fouchier (Rotterdam, The Netherlands), and A/HK/7/87 (A/H3N2 subtype) was obtained from Prof. J. Neyts (KU Leuven, Belgium).

### 5.4. Assessment of the antiviral activity of selected strains against HCoV-229E, RSV and influenza viruses in human cells

Bacterial strains were prepared as described above with UV-inactivation. The antiviral evaluation of UV-inactivated bacteria against HCoV-229E was performed by seeding Huh7 cells into 384-well plates. After 24h at 37°C, 5 fold serial dilutions of the bacteria (or PBS as a negative control) were added to the cells immediately prior to infection with HCoV-229E at 30 CCID_50_ (50% cell culture infective doses) per well. At 3 days post-infection, the virus-induced cytopathogenic effect was measured colorimetrically by the formazan-based 3-(4,5-dimethylthiazol-2-yl)-5-(3-carboxymethoxyphenyl)-2-(4-sulfophenyl)-2H-tetrazolium (MTS) cell viability assay (CellTiter 96 AQueous One Solution Cell Proliferation Assay from Promega, Madison, WI), and the antiviral activity was expressed as the 50% effective concentration (EC_50_). In parallel, the 50% cytotoxic concentration (CC_50_) was derived from mock-infected cells.

The evaluation of viral inhibition by UV-inactivated bacteria after pre-incubation with influenza viruses (A/H1N1 A/Ned/378/05, A/H3N2 A/HK/7/87 and B/Ned/537/05), respiratory syncytial virus and human coronavirus (HCoV-229E) was performed by seeding MDCK, HEp-2 or Huh7 cells into 384-well dishes, respectively, and the plates were incubated for 24h at 35°C (MDCK) or 37°C (HEp-2 and Huh7). One hour prior to infection, aliquots of UV-inactivated bacteria in PBS (or PBS alone as a negative control) were centrifuged and the pellets were resuspended in an equal volume of infection medium containing the viruses diluted to yield 30 CCID_50_ per well at infection. The mixtures were incubated at 37°C, 5% CO_2_ for 1h, and subsequently centrifuged (3000 g for 15 minutes at room temperature). The virus containing supernatants were used for infection of the respective virus host cells. At 3 days (HCoV-229E) or 4 days post-infection (influenza and RSV), the virus-induced cytopathogenic effect was measured colorimetrically by the formazan-based MTS assay, and the antiviral activity was expressed as the % viability relative to the uninfected control. In parallel, the % viability was measured in mock-infected MDCK, HEp-2 or Huh7 cells to assess possible cytotoxic effects.

### 5.5. Formulation of the microbiome throat spray and assessment of bacterial viability

The microbiome spray consisted of freeze-dried *L. casei* AMBR2, *L. plantarum* WCFS1, and *L. rhamnosus* GG, in a ratio of 50%, 33.3% and 16.7%, respectively. Viability of the powders from the individual strains was assessed at 4°C and 25°C every 4 weeks over a time period of 6 months via resuspension of the powders in PBS and plating out serial dilutions on MRS agar. The amount of CFU/g powder was evaluated compared to the start concentration. For the mixture of the powders or the final spray formulation, viability was assessed at 4°C, 15°C and 25°C over a period of 6 months. Every 4 weeks, powders (after suspension in PBS) or spray were plated out in serial dilutions on MRS agar to measure the amount of CFU/g powder or spray.

### 5.6. Evaluation of lactobacilli retention in the throat of healthy participants

To evaluate whether the bacteria in the spray are able to temporary colonize the throat, 12 healthy male and female adult participants were asked to use the spray and collect swabs of the throat at the start, after 30 minutes, and after 2 hours of spray administration. The study was conducted according to the guidelines of the Declaration of Helsinki, and approved by the Committee of Medical Ethics UZA/UAntwerpen (B3002021000018; approved 25^th^ of January 2021; ClinicalTrials.gov Identifier NCT04793997). Informed consent was obtained from all participants involved in this study prior to inclusion. Samples were registered and stored at the Biobank Antwerpen, Antwerp, Belgium; ID: BE 71030031000.

At each time point, 2 throat swabs were collected with eNAT™ swabs for microbial DNA extraction using the PowerFecal DNA isolation kit (Qiagen), and in PBS for lactobacilli cultivation on MRS agar. Dual-index paired-end sequencing of the throat samples was performed on the V4 region of the *16S rRNA* gene on a MiSeq Desktop sequencer (M00984, Illumina), as previously described (De Boeck et al. 2017,2019). For qPCR analysis, species-specific primers for *L. casei* AMBR2, *L. plantarum* WCFS1 and *L. rhamnosus* GG were designed (Table 2). Initially, a standard curve for each species was made to estimate the Ct∼CFU ratio. The expression of the genes was quantified by RT-qPCR on a StepOne Plus Real-Time PCR System (v.2.0; Applied Biosystems, Foster City, California, United States). Each DNA sample was amplified with PowerSYBR® Green PCR Master Mix (Applied Biosystems) in a total volume of 20 μL with 0.15 μM of each primer, 40 ng of cDNA and nuclease-free water.

**Table 2.**
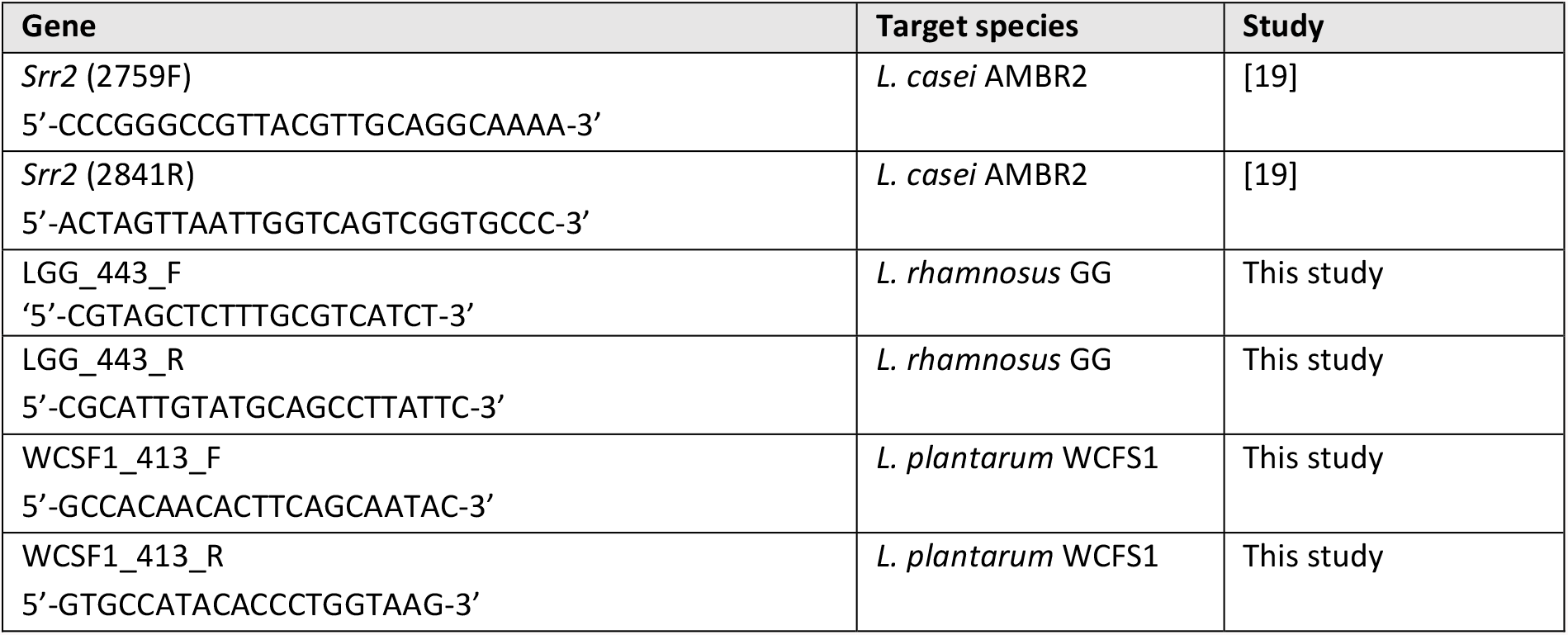
qPCR primers used in this study.

Throat swabs were collected by swabbing along the back of the throat and both tonsils and cultivated after resuspension in 1 ml PBS and plating out serial dilutions on MRS agar. Plates were incubated for 2 days at 37°C. Single colonies were identified with PCR and Sanger sequencing of the full *16S rRNA* gene using the 27F (5’-AGA GTT TGA TCM TGG CTC AG-3’) and 1492R (5’-GGT TAC CTT GTT ACG ACT T-3’) primers.

### 5.7. Statistical data analysis

Data from *in vitro* assays was analysed in GraphPad Prism version 9.2.0. Normality testing was performed on data from *in vitro* assays with bacteria using the Kolmogorov-Smirnov test followed by One-way ANOVA with Dunnett’s multiple comparisons test, or the Kruskal-Wallis test with Dunn’s multiple comparisons test. Differences were considered significant at p < 0.05.

### 5.8. Data availability statement

The datasets generated and analysed during the current study are available in the European Nucleotide Archive (ENA) under accession number PRJEB49183 after publication of this manuscript.

## 6. Declarations

### 6.3. Author Contributions

Conceptualization, SL, IDB, IS, IC, TH, PAB, PD, SC, VV, MvdB; methodology, IDB, IS, SL, LP, IC, TH, AS, IG, LD, EC; validation, IDB, IS, EC, LD, TH, AS, IG; formal analysis, IDB, IS, EC, LP, AS; investigation, IDB, IS, EC, LD, LP, TH, AS, PAB, SL, IG, LP; resources, SL, DS, IC, PD; data curation, IDB, IS, EC, TH, IC; writing— original draft preparation, IDB, IS, SL; writing—review and editing, all authors; visualization, IDB, IS, EC, AS; project administration, IDB, IS, SL, TH, IC; funding acquisition, SL, DS, IC. All authors have read and agreed to the published version of the manuscript.

## 6.4. Acknowledgments

The authors want to thank the entire research group of the Lab of Applied Microbiology and Biotechnology of the University of Antwerp. We would also like to thank Kate Van Loock and Lorenzo Carreon for the help with the formulation development and the Biobank Antwerpen for the storage and registration of samples of human origin.

